# Winner-Takes-All Resource Competition Redirects Cascading Cell Fate Transitions

**DOI:** 10.1101/2020.05.23.103259

**Authors:** Rong Zhang, Hanah Goetz, Juan Melendez-Alvarez, Jiao Li, Tian Ding, Xiao Wang, Xiao-Jun Tian

## Abstract

Failure of modularity remains a significant challenge for synthetic gene circuits assembled with tested modules as they often do not function as expected. Competition over shared limited gene expression resources is a crucial underlying reason. Here, we first built a synthetic cascading bistable switches (Syn-CBS) circuit in a single strain with two coupled self-activation modules to achieve two successive cell fate transitions. Interestingly, we found that the *in vivo* transition path was redirected as the activation of one switch always prevailed against the other instead of the theoretically expected coactivation. This qualitatively different type of resource competition between the two modules follows a ‘winner-takes-all’ rule, where the winner is determined by the relative connection strength between the modules. To decouple the resource competition, we constructed a two-strain circuit, which achieved successive activation and stable coactivation of the two switches. We unveiled a nonlinear resource competition within synthetic gene circuits and provided a division of labor strategy to minimize unfavorable effects.

## INTRODUCTION

Modularity is one of the important design principles for engineering sophisticated synthetic gene circuits by breaking the system down into small modules to reduce complexity. However, the whole circuit often does not function as expected when the tested modules are assembled even after several rounds of design-build-test iterations. One of the most important reasons is that various hidden circuit-host interactions, including resource competition, could significantly perturb the performance of synthetic gene circuits ^1–4^. The available cellular resources in the host cell, such as transcriptional and translational machinery (e.g., RNA polymerases and ribosomes), are limited for synthetic gene circuits, thus resulting in undesired competition between the modules within one gene circuit. For example, resource competition causes retroactivity from downstream regulators to upstream dynamics and alters the expected behaviors^5,6^. Thus, it is essential to characterize how the modules in one circuit are unintentionally coupled because of the limited amount of shared resources and how this coupling leads to the loss of modularity.

The coupling between two separated genes in the same plasmid is found to be constrained by an inverse linear relationship, analogous to the isocost lines used in economics ^7,8^. The dependence between genetic loads and gene expression is also found to be governed by equations analogous to Ohm’s law used in electrical circuits ^9^. These elegant equations indeed help us to quantitatively understand the resource competition between simple circuits. However, the behavior between more complex modules within one gene circuit, such as positive feedback loops, remains unclear.

Here, in this work, we built a synthetic cascading bistable switches (Syn-CBS) circuit to achieve successive cell fate transitions. First, we constructed a single strain Syn-CBS circuit with two mutually connected self-activation modules to achieve stepwise activation of two bistable switches by controlling the inducer dose. Interestingly, we found that an increase of the inducer changes the system from a state in which only one of the switches is activated to a state in which the other switch is the only one activated, instead of the theoretically expected coactivation of both switches. The underlying reason was that the two modules may be competing for limited resources and thus inhibit each other indirectly. This winner-takes-all (WTA) behavior resulting from resource competition between the two connected modules was also verified in controlled experiments when the modules were disconnected. We further found the relative strength of connections between the modules determines the winner in the syn-CBS circuit. To decouple the resource competition between the two modules, we constructed a two-strain Syn-CBS circuit that could achieve stable coactivation of the two coupled bistable switches. Thus, the effect of the resource competition on the Syn-CBS circuit is minimized through a division of labor using microbial consortia.

## RESULTS

### Design of Syn-CBS circuit to achieve successive cell fate transitions

The existence of multiple stable states under the same condition, also known as multistability, plays a critical role in diverse biological processes ^10–18^. Previously, we mathematically predicted and experimentally verified that epithelial-to-mesenchymal transition (EMT) is a two-step process governed by cascading bistable switches (CBS) ^14,15^. To further understand the design principle of CBS for achieving successive cell fate transitions, we designed a synthetic CBS (Syn-CBS) circuit with two mutually regulated modules to study the design principle of cascading cell fate transitions. In this design (Fig. 1A), one module (M1) is designed with self-activation of AraC, which is controlled by Arabinose (L-ara), and the other (M2) is designed with self-activation of LuxR and is controlled by quorum-sensing signal 3oxo-C6-HSL (C6). GFP with LVA tag (GFP-lva) and RFP with AAV tag (RFP-aav) are used as the outputs of the two switches, respectively. We have tested that both individual modules could function as a bistable switch ^3^. Thus, with correct connections of the two modules, we expected to see the successive activation of these two bistable switches. That is, with increase in the dose of the inducer L-ara, the Syn-CBS strain should transition from a state with no activation of either switch to a state with only one switch activated, and then to a final state with both switches activated.

**Figure 1.**
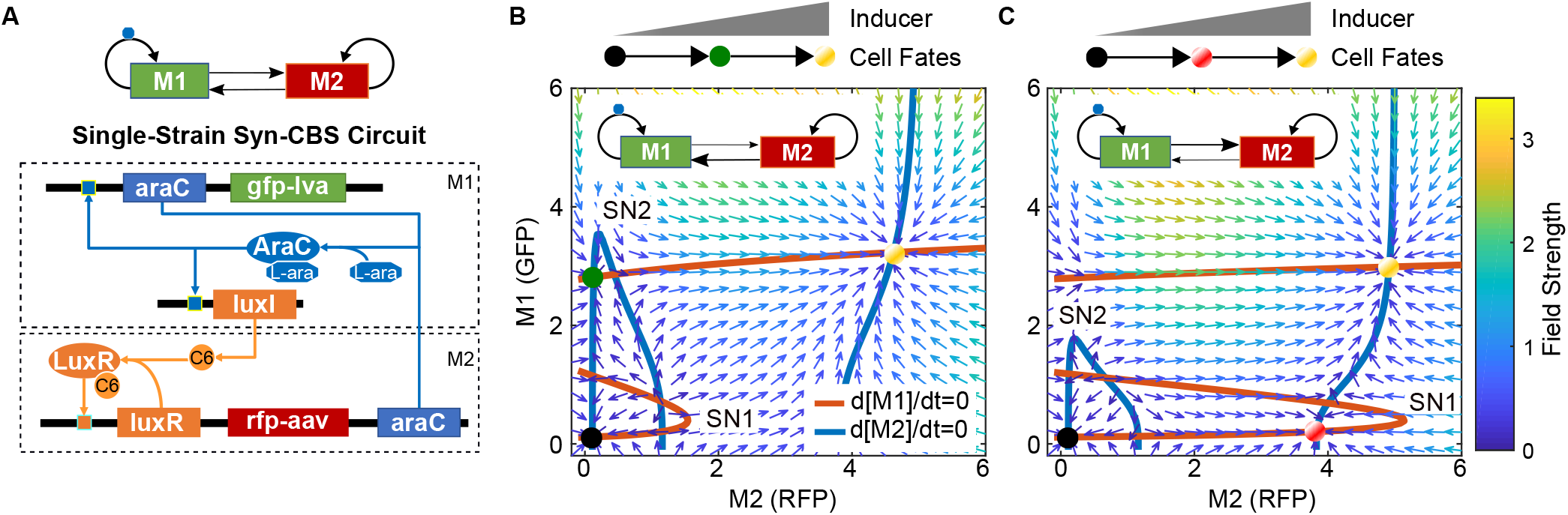
Conceptual design of the synthetic cascading bistable switches (Syn-CBS) circuit. (A) Diagram of the Syn-CBS circuit, in which two self-activation modules mutually activate each other. The araC self-activation in module 1 (M1), regulated by L-ara, is designed to achieve one bistable switch, and the luxR self-activation in module 2 (M2), regulated by C6, is designed to achieve another bistable switch. (B-C) Phase plane analysis shows the two different expected cell fate transition paths depending on the strength of the links between the two switches. (B) A weak M1-to-M2 link and strong M2-to-M1 link leads to a cell fate transition from a RFP-low/GFP-low state, to a RFP-low /GFP-high state, and then to a RFP-high/GFP-high state. (C) A strong M1-to-M2 link and weak M2-to-M1 link leads to a cell fate transition from a RFP-low/GFP-low state, to a RFP-high/GFP-low state, and then to a RFP-high/GFP-high state. The nullclines of M1 and M2 are shown in red and blue, respectively. The vector field of the system is represented by small arrows, where the color is proportional to the field strength. The three cell fates are indicated by filled circles at the intersections of the two nullclines.

To demonstrate that this circuit design could achieve successive cell fate transitions, we developed a mathematical model for the Syn-CBS circuit (see SI for details). Through graphical analysis of the nullclines, vector field, and potential landscape in the M1-M2 phase plane, the model predicted that this system could achieve a stepwise activation of the two switches in two different ways (Fig. 1BC, S1). The nullclines analysis shows that both the M1 and M2 modules could function as a bistable switch with the other as an input (Fig. 1BC). That is, both modules need the other to be above a certain threshold for activation (SN1 for M1 and SN2 for M2). However, the activation thresholds depend on the strengths of the two links between the two modules. If the strength of the M2-to-M1 connection is strong and the M1-to-M2 connection is weak, the dose of L-ara required for M1 switch activation is smaller than the dose needed for M2 switch activation. That is, the threshold SN1 is smaller than SN2 (Fig. 1B). The corresponding nullcline intersections give three stable steady-states: LL (low-RFP/low-GFP), LH (low-RFP/high-GFP), and HH (high-RFP/high-GFP). With increase of the L-ara dose, both the levels of M1 and M2 increase. However, the M1 switch is activated first, turning the cells green due to the low activation threshold. The M2 module is activated later, turning cells yellow under a larger dose of L-ara. The quasi-potential landscape was calculated by solving the corresponding Chemical Master Equation (CME) (see SI for details) to visualize the three cell fates as potential wells (Fig. S1A). A design with a weak M2-to-M1 connection and a strong M1-to-M2 connection leads to a flipped scenario, in which cells transition from a LL state to a HL state and then to a HH state (Fig. 1C, S1B). Thus, the Syn-CBS circuit is a good theoretical design to achieve successive cell fate transitions.

### Resource competition deviates cell fate transition from the desired stepwise manner

Next, we constructed and put the whole Syn-CBS circuit in *E. Coli* strain. We first studied the relationship between the two modules by measuring the mean GFP and RFP levels using a plate reader, and found that GFP vs. RFP showed a negative relationship (Fig. 2A), instead of the positive one expected from theoretical design, suggesting resource competition between the two modules. Interestingly, this inverse relationship followed a two-phase piecewise linear function, showing a shallow slope when the GFP level was high (black curve, Fig. 2A) and a steep slope when the RFP level was high (red curve, Fig. 2A). To further study this phenomenon, we measured the cell fate transition at the single-cell level with flow cytometry under various concentrations of L-ara. Unexpectedly, three different separate stable steady-states were found in the GFP/RFP space (Fig. 2B). With increase of the inducer L-ara, most of the cells first entered a high-RFP/low-GFP state, then, to our surprise, jumped to a low-RFP/high-GFP state with only a few cells staying in the high-RFP/high-GFP state (Fig. 2B, S2). Thus, experimental data showed a slim chance for the existence of the theoretically expected coactivation high-RFP/high-GFP state, which we now consider as an unstable state. That is, the desired path of cell fate transitions was redirected from the HH state to the LH state (Fig. 2C). The two moduels competed for limited resources, thus inhibited each other indirectly (Fig. 2D), an idea not included in the Syn-CBS gene circuit design.

**Figure 2.**
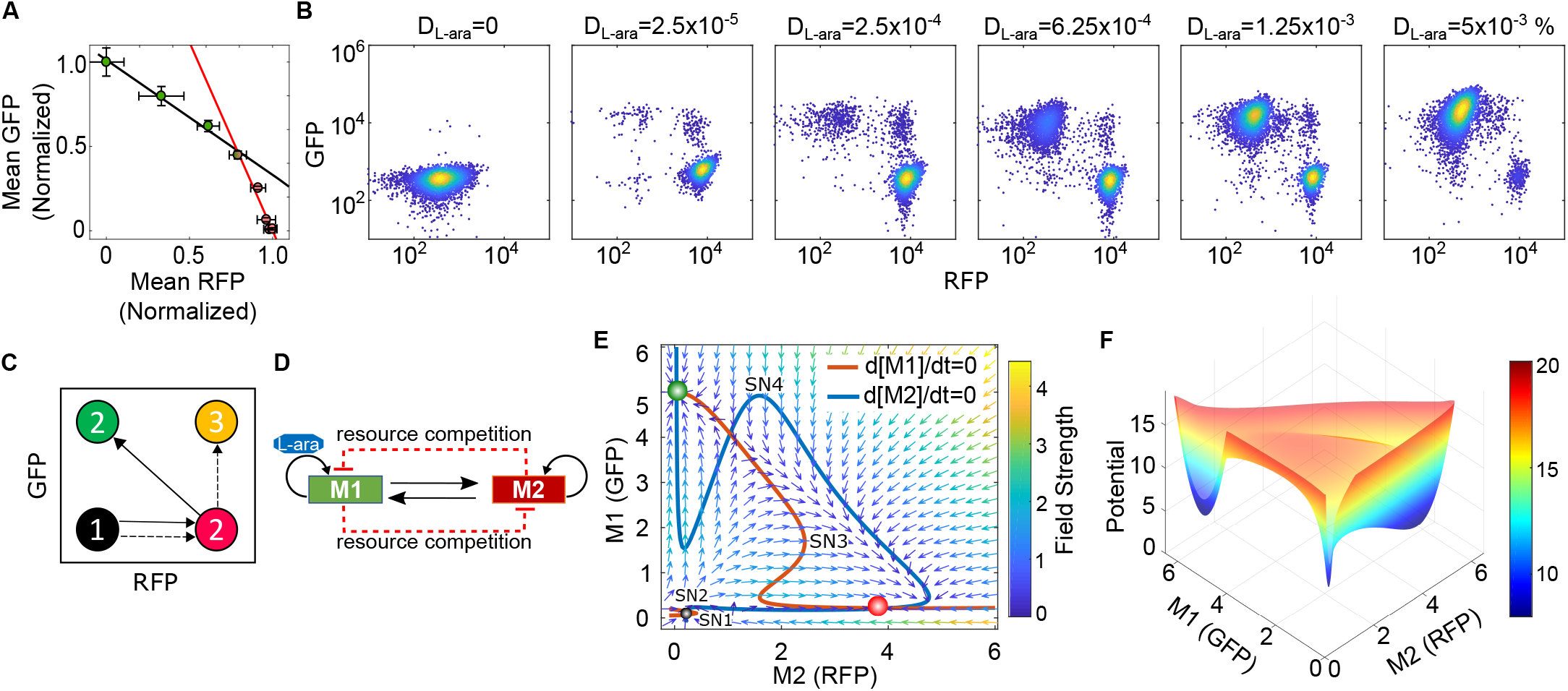
Resource competition deviates the cell fate transitions in the one-strain Syn-CBS circuit. (A) The normalized steady-state of average GFP vs. RFP measured by plate reader shows a two-phase piecewise linear relationship. Data displayed as mean ± s.d. (n = 3). (B) Flow cytometry data shows cell state transitions in one-strain Syn-CBS circuit with increasing level of inducer L-ara (D_L-ara_). 10,000 events were recorded for each sample by flow cytometry. (C) Diagram of the perturbed state transitions by resource competition. Dash line: expected path. Solid line: perturbed path. (D) Diagram of the revised model by including resource competition. (E) Phase plane diagrams. The nullclines of M1 and M2 are shown in red and blue, respectively. The vector field of the system is represented by small arrows, where the color is proportional to the field strength. The three cell fates are indicated by filled circles at the intersections of the two nullclines. (F) Calculated potential landscape.

The discrepancy between the model prediction and experimental data was resolved by including resource competition into the model (see SI for details). The shapes of the new nullclines changes in comparison to the Syn-CBS model without resource competition (Fig. 2E and 1B, respectively). The behavior is similar in the sense that M1 activation needs M2 above a certain threshold (SN1 on the red curve, Fig. 2E). Nonetheless, the continuous increase of M2 now turns the M1 switch off after going above yet another threshold (SN3 on the red curve, Fig. 2E). Additionally, the M2 nullcline is changed to reflect that M2 can switch off as M1 increases (SN4 of the blue curve, Fig. 2E). Thus, the intersections of the two nullclines give three different steady states: the LL state (black circle), LH state (green circle), and HL state (red circle). The quasi-potential landscape also shows three potential wells corresponding to three cell fates without coactivation (Fig. 2F). Although the system has a potential to be quadrastable, the strong mutual inhibition from indirect resource competition prevents the nullclines from intersecting in the high-RFP/high-GFP state, as was suggested by our experimental data. Taken together, our results suggest that resource competition between the two modules in one gene circuit deviates cell fate transitions from the desired stepwise manner.

### WTA Behavior Found in the Resource Competition Between Separated Bistable Switches

To further confirm the resource competition between the two modules in the Syn-CBS circuit, we studied the behaviors of the two separated bistable switches (Syn-SBS) system, in which the previous links between the two modules in the Syn-CBS circuit were removed (Fig. 3A). The network topology (Fig. 3A, right) is similar to the synthetic circuit MINPA ^13^ and cell differentiation system ^19–22^, with the difference being a mutual inhibition mediated by resource competition. The cell fate transition was induced by increasing doses of C6 combined with a fixed dose of L-ara added at the beginning of the experiment and measured with flow cytometry (Fig. 3B). It is noted that the activation speed of the bistable switch elevates while the dose of its inducer increases^3,23^. Thus, under low dose of C6, M1 is activated so fast that M2 is repressed from activation. Under a moderate dose of C6, the activation speed of the two modules are similar, thus leading to their coactivation. However, a high dose of C6 activated the M2 switch so fast that M1 was completely blocked by M2 for activation (Fig.3B, S3). This results in a ‘winner-takes-all’ (WTA) behavior with the presence of resource competition between the two modules, i.e., the first activated switch always suppresses the activation of the second switch.

**Figure 3.**
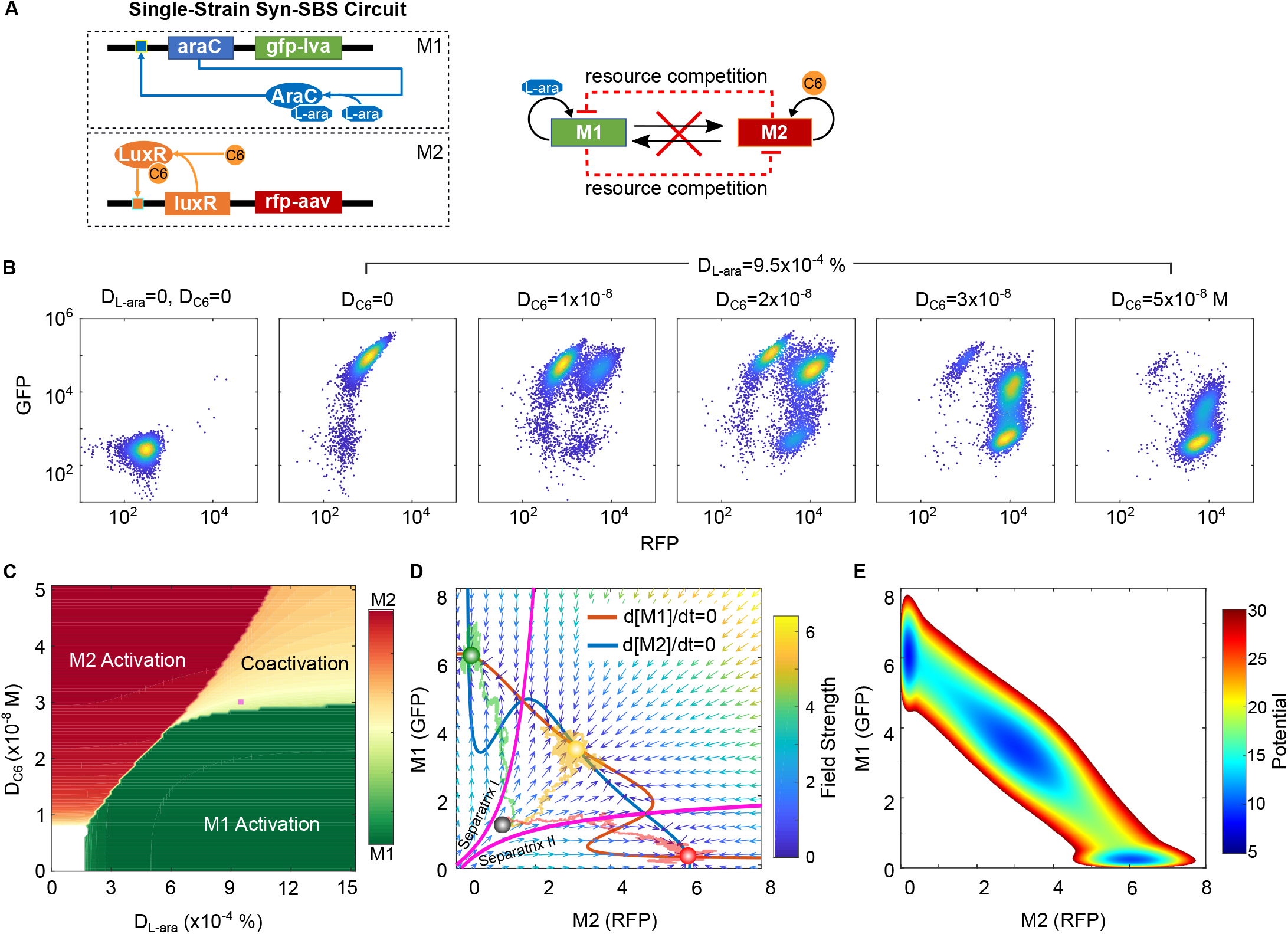
Resource competition between two separate bistable switches. (A). Diagram of the two separate bistable switches (Syn-SBS). (B). Flow cytometry data shows cell state transitions with an increasing level of inducer C6 (D_C6_) and a fixed dose of L-ara (D_Lara_=9.5*10^−4^%). 10,000 events were recorded for each sample by flow cytometry. The two inducers were both added at 0 hr. (C). Cell fates in the space of two inducers L-ara and C6. (D). Simulated stochastic trajectories in the phase plane diagrams. The nullclines of M1 and M2 are shown in red and blue, respectively, while separatrices are shown in pink. The vector field of the system is represented by small arrows, where the color is proportional to the field strength. The three cell fates (red, green, and yellow circles) are found at the intersections of the two nullclines. Three representative single-cell stochastic trajectories (green, yellow, and red trails) show the evolution of the system from the same initial condition (black circle, D_Lara_ = 0 % and D_C6_=0 M) to three different states with the same induction (D_Lara_ = 6.5×10^−4^ % and D_C6_=0.60 M). (E) Calculated potential landscape.

In order to understand the mechanisms of the WTA phenomena, we conducted simulation with a mathematical model for the Syn-SBS system (see SI for more details). As shown in Fig. 3C, the simulated cell fates can be represented by the levels of M1 and M2 in the dose space of two inducers. The space is divided into four regions: no switch activation at the corner with low C6 and low L-ara, M2 switch activation only at the corner with high C6 and low L-ara, M1 switch activation only at the corner with low C6 and high L-ara, and coactivation of the two switches at the corner where the two inducers are high and well-balanced. In Fig. 3D, three stochastically simulated single-cell trajectories with a fixed dose of L-ara and C6 were shown in the M1-M2 phase plane and direction field. Three stable steady-states, including two states with only one winner and the coactivation state, can be found at the intersections of the nullclines (red and blue curves, Fig. 3D) and the potential landscape (Fig. 3E). The existence of the coactivation state in the Syn-SBS system may result from a smaller burden compared to that of the Syn-CBS circuit, given that two more genes are included in the latter. It is noted that if one cell crosses Separatrix I (uppermost pink curve in Fig. 3D), M2 starts to be repressed by M1 due to the resource competition, leading the commitment of the cell to the GFP-high state. Similarly, the cell becomes committed to the RFP-high state after crossing Separatrix II (lowermost pink curve in Fig. 3D). If the levels of M1 and M2 are well-balanced, the cell is able to reach the coactivation state between the two separatrices.

To fully comprehend how sequential activation of the two bistable switches in the one strain Syn-CBS circuit failed due to resource competition, we then studied how sequential addition of the two inducers affects the cell fate transitions in the Syn-SBS system. We fixed the doses of both inducers, adding L-ara initially but varying the addition time point of C6 from 0 to 4 hours, to mimic the design of the Syn-CBS circuit (Fig. S4A). As shown in Fig S4B, when both inducers were added at time point 0, the M2 switch won the competition, seen by the fact that most of the cells are in the RFP-high state, which was consistent with Fig. 3B. However, when C6 was added 1-2 hours later, more and more cells show coactivation of both switches (Fig. S4BC). When C6 was added 4 hours later, most of the cells show only high GFP (Fig. S4BC). That is, the M1 switch was activated first and started to repress the activation of the M2 switch by taking all the available resources. It is noted that the inactivation of the M2 switch was not due to the late addition of C6 since the M2 switch was able to be activated in the parallel experiment with the same C6 dose and time points but without L-ara (Fig. S4B, bottom). In addition, the simulated cell fates with a fixed L-ara dose are shown in the space of D_C6_ and T_C6_ (Fig. S4D): M1 switch activation only (in the region with low C6 or late C6 addition), M2 switch activation only (in the region with high C6 and early addition), and coactivation of the two switches (in the region with moderate C6 and early addition). Representative stochastic single-cell trajectories with three C6 addition time points were shown in the M1-M2 phase planes (Fig. S4E). The trajectories of the cells were first following the direction filed in the M1-M2 phase plane without C6 to the GFP-high state before C6 addtition(left panel), and then following the direction filed in the phase plane with C6 to three different states after C6 addition (right panel). At the time point after the cell has crossed Separatrix I (leftmost pink curve, right panel of Fig. S4E), adding C6 does not activate the M2 switch (green trajectory). At the time point where the cell has not yet crossed Separatrix I, adding C6 may lead to coactivation (yellow trajectory). Adding C6 early on may lead to the cell crossing Separatrix II (lowermost pink curve, right panel of Fig. S4E) and M2 activation only (red trajectory). Taken together, these results confirmed the WTA behavior when resource competition was present between the two modules.

### Relative strength of module connections determines the winner of resource competition

To further understand how the winner was determined during the resource competition between the modules within the Syn-CBS circuit, we studied if the strength of the module connections affected the outcomes of the cell fate transitions by finetuning the M1-to-M2 link. A hybrid promoter Para/tet was used to control the production of C6 to tune the module connection by external chemical inducer anhydrotetracycline hydrochloride (aTc). As shown in Fig. 4A, luxI gene expression is jointly regulated by AraC and tetR, while tetR is negatively controlled by aTc.

**Figure 4.**
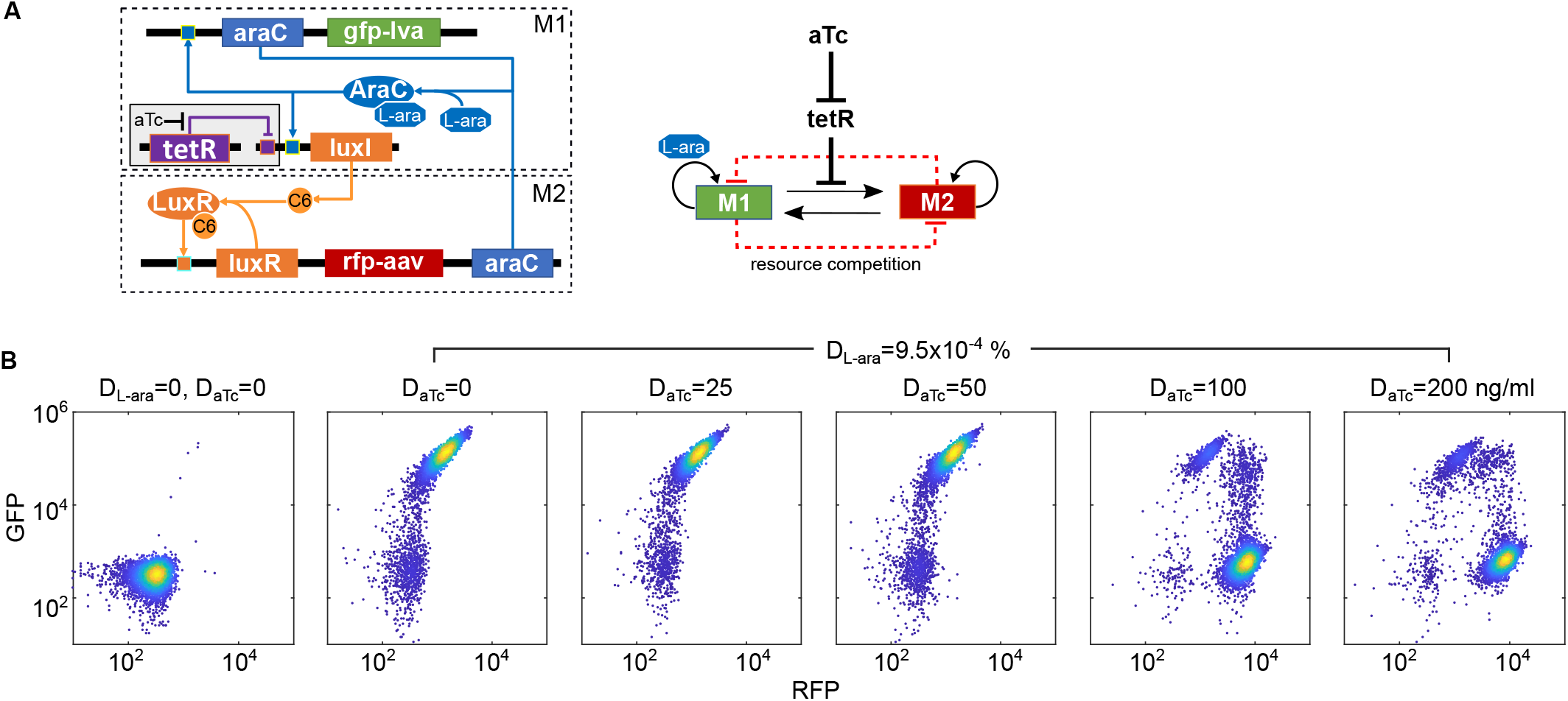
Relative strength of module connections determines the winner of resource competition. (A) Diagram of the Syn-CBS circuit with a tetR module for fine-tuning the connection between two bistable switch modules. A hybrid promoter Para/tet is used for controlling the production of C6 to tune the connection between the two switches. (B) Flow cytometry data shows cell state transitions with various doses of inducer aTc (D_aTc_) and a fixed dose of L-ara (D_L-ara_). 10,000 events were recorded for each sample by flow cytometry.

With this design, we used a high enough dose of L-ara so that the M1 switch could activate. We then increased the dose of aTc to release the inhibition of C6 production by tetR and activate the M2 switch. As shown in Fig. 4B, the M1 switch was activated without aTc in the presence of L-ara. An increase in the dose of aTc did activate the M2 switch as expected, but the M1 switch was then blocked from activation (Fig. 4B and Fig. S5A). These results suggest that the relative strength of the module connections determines the winner of the resource competition in the Syn-CSB circuit. The simulated cell fates in the space of L-ara and aTc shows that the M1 switch can only be activated with high L-ara and low aTc, while the M2 switch is only activated with high L-ara and high aTc (Fig. S5B), which is consistent with the experimental data. Taken together, although the two bistable switch modules in the Syn-CBS circuit are designed to be mutually activated, they race against each other for the limited resources in order to activate. That is, the first activated module takes available resources and thus inhibits the capability of the other to switch on. Since the Syn-CBS circuit was designed to achieve the sequential activation of the two switches, the use of only one strain makes the WTA behavior and failure in the modularity design of the circuit inevitable. Therefore, we need to decouple these indirected hidden linkes between the modules.

### Stabilized coactivation of the two switches through a division of labor using microbial consortia

In order to decouple the undesired crosstalk within the gene circuit due to resource competition, we designed and constructed two-strain Syn-CBS circuits by dividing the two modules into two separate cells (Fig. 5A), instead of placing one whole gene circuit in a single cell. We considered these new two-strain Syn-CBS circuits both with and without the tetR module and kept the circuit connections similar to that of the one-strain Syn-CSB circuits (Fig. 1A, 4A).

**Figure 5.**
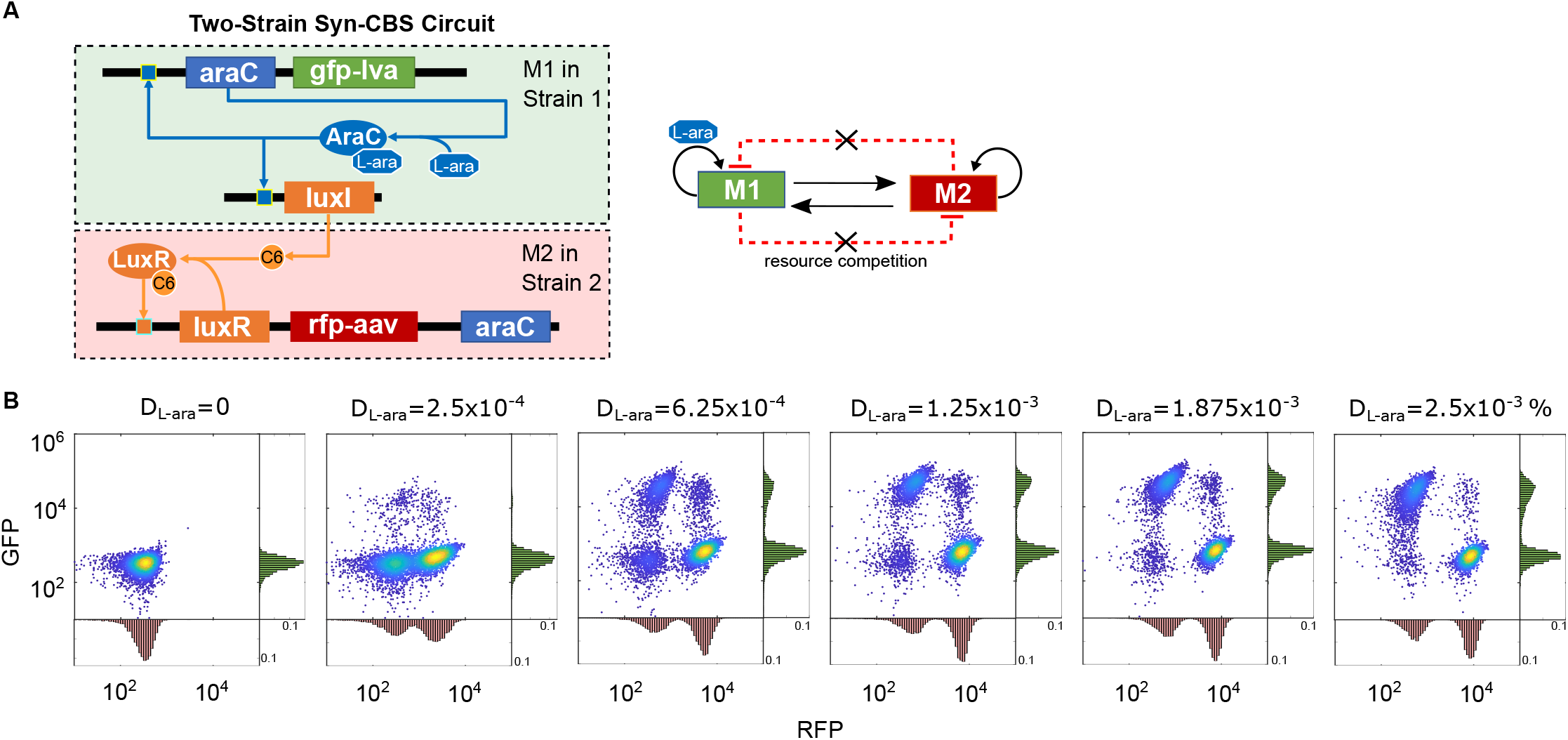
Minimize resource competition through a division of labor using microbial consortia. (A) Diagram of two-strain Syn-CBS circuits without a tetR module. (B) Flow cytometry data shows the expected stepwise cell state transitions by increasing the dose of inducer L-ara (D_Lara_). 10,000 events were recorded for each sample by flow cytometry.

We systematically studied the cell fate transitions with the two-strain Syn-CBS circuits. For the design without the tetR module, a low dose of L-ara was enough to transition some cells into a high-RFP state (Fig. 5B). As the dose of L-ara increased, the rest of the cells gradually transitioned to a high-GFP state and became stable under a high dose of inducer (Fig. 5B, S6). It is noted that GFP and RFP were in different strains, so stable coactivation of the two modules is instead represted by the RFP^hi^/GFP^lo^ and RFP^lo^/GFP^hi^ populations (Fig. 5B, S6A). Similarly, in the two-strain Syn-CBS circuit with the tetR module (Fig. S7A), as the aTc dose risen, part of the cells first transitioned to a GFP-high state under a low aTc dose, while the rest of the cells continuously transitioned to a RFP-high state (Fig. S7B-C). The stable coexistence of the two populations was also found. It is noted that due to the heterogeneity of the growth rates of the two strains, the fractions of the cell populations were not well-balanced, but the overall ratio did not change over a dose range of the inducer in both circuits. In addition, the weak anticorrelation between the two switches in both two-strain Syn-CBS circuits when under a high dose of inducer suggests the adverse effects of resource competition are minimized through a division of labor. Thus, the two-strain Syn-CBS circuits work better to achieve the successive activation of the two bistable switches without the result of one switching off.

## DISCUSSION

Resource competition is commonplace at various levels of regulation in biological systems, including transcriptional, translational, and post-translational. Resource competition can be exploited to its best advantage for natural and synthetic biological systems. For example, amplified sensitivity arises from covalent modifications with limited enzymes and molecular titration ^24–28^. Competition for limited proteases was utilized to coordinate genetic oscillators ^29^. Adding competing transcriptional binding sites on sponge plasmid makes the repressilator more robust ^30^. However, resource competition within one gene circuit may also change circuit behaviors ^5,6^. Here, we showed that it was challenging to achieve successive activation of two bistable switches in one strain due to resource competition. We also found that the two bistable switches competed for limited resources and that this leads to only one winner that takes all the available resources. Interestingly, the outcomes of the WTA competition depended on the dynamics of the two switches, given that the faster one was always the winner.

Several approaches have been proposed to counteract the effects of resource competition, either by finetuning the parameters in the gene circuit ^5,7^ or manipulating the size of the orthogonal resource pools ^31–34^. Additionally, a burden-driven negative feedback loop was implemented to control gene expression by monitoring the cellular burden ^35–37^. The negative feedback loop can also be integrated within synthetic gene circuits to control resource competition ^8,38–40^. In this study, we compared the single- and two-strain Syn-CBS circuits and found that the deviated cell fate transitions due to resource competition in monoclonal microbial are corrected in micro-organism consortia. A trade-off was found between robustness to environmental disturbances and robustness to perturbations in available resources for the genetic circuit ^41^. Synthetic microbial consortia have been used for engineering multicellular synthetic systems ^42–44^ and metabolic pathways ^45^.

Resource competition also exists between the host cell and the synthetic gene circuit. Thus, the strategies of the host cells on resource allocation also influence the performance of the gene circuits. Host cells are dynamically adjusting their intracellular resource reallocations in response to nutrient availability or shifts ^46–48^. Therefore, the availability of cellular resources to the synthetic gene circuits is also very dynamic and stochastic. Recently, it was found that bacterial strategies differ in their response to starvation for carbon, nitrogen, or phosphate ^49,50^. It is very challenging to accurately predict the circuit behaviors under the conditions of dynamic resource allocation. An integrative circuit-host modeling framework has been developed to predict behaviors of synthetic gene circuits ^4^. Dynamic models of resource allocation were also developed in response to the presence of a synthetic circuit ^51,52^.

Our recent work found that synthetic switches may lose memory due to cell growth feedback depending on their network topology ^3^. We mathematically and experimentally demonstrated that the self-activation gene circuit is very sensitive to the growth feedback. In contrast, the toggle switch circuit is very robust, although the gene expression of both circuits was decreased significantly due to the fast cell growth ^3^. Recently, McBride et al. mathematically proved that the mutual activation circuit and reciprocal inhibition circuit also behave differently under the context of resource competition ^53^. Similarly, the repression cascade seems more robust in contrast with the activation cascade ^5^. All of these works suggest that the perturbation of the circuit function depends on the network topology and thus the context of various circuit-host interactions needs to be considered for gene circuits design.

## MATERIAL and METHODS

### Strains, media, and chemicals

E. coli DH10B (Invitrogen, USA) was used for all the cloning constructions. Measurement of the circuits was performed in *E. Coli* K-12 MG1655ΔlacIΔaraCBAD as described in ^3^. Cells were grown in 15 ml tubes with 250 rotations per minute at 37 °C in Luria-Bertani broth (LB broth) with 100 μg/ml chloramphenicol or 50 μg/ml kanamycin. L-ara (L-(+)-Arabinose, Sigma-Aldrich) C6 (3oxo-C6-HSL, Sigma-Aldrich) and aTc (Anhydrotetracycline hydrochloride, Abcam) were dissolved in ddH2O and later diluted to appropriate working solutions. All the oligos DNA were synthesized by Integrated DNA Technologies, Inc. (IDT).

### Plasmids construction

The generation of araC gene and promoter Plux was described in ^3^. The sequence and characterization of luxRG2C can be found in ^54^. Two sets of primers were used to generate luxRG2C sequence from template BioBrick C0062. Primer set one was: forward 5’-ctggaattcgcggccgcttctagatgaaaaacataaatgccgac-3’ and reverse 5’-ggactgcagcggccgctactagtagtttattaatttttaaagtatgggcaatc-3’; primer set two was: forward 5’-gtttagtttccctattcatacggctaacaatggcttcggaatacttagttttgcacattc-3’ and reverse 5’-gtatgaatagggaaactaaacccagtgataagacctgctgttttcgcttctttaattac-3’. The gene sequence of unstable RFP tagged with AANDENYAAAV ^55^ peptide tail (RfpAAV) was synthesized by PCR using BioBrick K1399001 as a template and primer set: forward 5’-tgccacctgacgtctaagaa-3’ and reverse 5’-gctactagtattattaaactgctgctgcgtagttttcgtcgtttgcagc-3’. The sequence of Para/tet is 5’-GCTTCTAGAGacattgattatttgcacggcgtcacactttgctatgccatagcaagatagtccataagattagcggatcctacctgacgctttttatcgcaactctctactgtttctccattccctatcagtgatagaTACTAGTAGCGGCCGCTGC AGTCC-3’, in which the lowercase part stands for the sequence for the promoter and the uppercase parts stands for the sequences flank the promoter which can be cut by restriction enzymes XbaI and PstI. All the modified parts were flanked by RFC 10 sequence from iGEM in order for them to be constructed the same way as standard BioBricks. The BioBricks used directly to build our circuits were listed in Table S1. All parts were first restriction digested using desired combinations of FastDigest restriction enzyme chosen from EcoRI, XbaI, SpeI, and PstI (Thermo Fisher) and separated by gel electrophoresis, and then purified using GelElute Gel Extraction Kit (Sigma-Aldrich) followed by ligation using T4 DNA ligase (New England BioLabs). Then the ligation products were transformed into E. coli strain DH10B and later the positive colonies were screened. Finally, the plasmids were extracted using GenElute Plasmids Miniprep Kit (Sigma-Aldrich). Each operon constituting the circuits was constructed monocistronically and its sequence was verified before combined into circuits. Details of all the operons and the circuits can be found in Table S2 and Table S3.

### Flow cytometry

All samples were analyzed using Accuri C6 flow cytometer (Becton Dickinson) with excitation/emission filters 480nm/530nm (FL1-A) for GFP detection and FL3-A for RFP at indicated time points. 10,000 events were recorded for each sample. At least three replicated tests were performed for each experiment. Data files were analyzed with MATLAB (MathWorks). Non-cellular small particles with FSA values lower than 30,000 were eliminated during data analysis according to data from the plain LB medium without any cells as a negative control. Some of the doublets in the data were eliminated with the gating in the space of FSC_A/FSC_H. This gating method was not able to remove the doublets of the small cells, and thus a tiny fraction of cells still can be seen in the figures.

### Circuit inductions

On day one, the plasmids carrying the circuits were inoculated into E. coli strain K-12 MG1655ΔlacIΔaraCBAD and were grown on LB plates with 50 μg/ml kanamycin overnight at 37 °C. On day two in the morning, one colony was picked and inoculated into 400 μl LB medium with 50 μg/ml kanamycin and was grown to OD about 1.0 (measured in 200μl volume by plate reader for absorbance at 600 nm) in a 5 ml culture tube in the shaker. The cells were then diluted 1000 folds into fresh LB broth with 50 μg/ml kanamycin and each portion of a 2 ml aliquot was distributed into 15 ml culture. Later, respective inducers were added into each tube, and the cells were grown for 16 hours in the shaker, and then data were gathered on flow cytometry.

### Average fluorescence analysis performed by Plate Reader

Synergy H1 Hybrid Reader from BioTek was used to perform the **average** fluorescence **analysis**. 200 μl of culture was loaded into each well of the 96-well plate. LB broth without cells was used as a blank. The plate was incubated at 37°C with orbital shaking at the frequency of 807 cpm (circles per minute). OD (optical density) of the culture was measured by absorbance at 600 nm; GFP was detected by excitation/emission at 485/515 nm.

### Mathematical models

Ordinary differential equation models were developed to describe and analyze all the synthetic gene circuits with or without considering resource competition at the population level. The stochastic simulation algorithm was developed to characterize the stochasticity at the single-cell level. The Chemical Master equation (CME) was used to calculate the steady probability distribution and estimate the potential landscape. Details are provided in the Supplementary Note.

## Acknowledgments

We thank Dr. XX for valuable comments. This project was supported by the ASU School of Biological and Health Systems Engineering and NSF grant (no. EF-1921412 to X-J.T.). H.G. and J.M.-A. were also supported by the Arizona State University Dean’s Fellowship.

## Author contributions

R.Z. and X.-J.T. conceived the study. X.-J.T. and R.Z. designed the study. R.Z. and J. L. performed experiments. J.M.-A., H.G., and X.-J.T. performed theoretical studies. J.M.-A., H.G., R.Z., D.T., X.W., and X.-J.T. analyzed the data. X.-J.T. wrote the manuscript. H.G., R.Z., X.W., and J.M.-A., edited the manuscript.

