## Supplementary Information for "Winner-Takes-All Resource Competition Redirects Cascading Cell Fate Transitions"

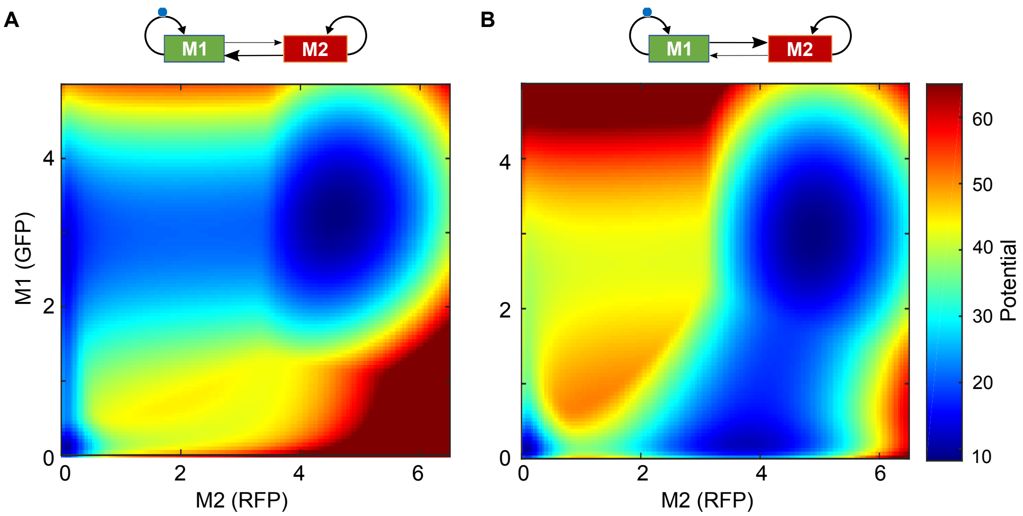


**Figure S1. Calculated potential landscape of the Syn-CBS circuit**. The mathematical model without resource competition was used here to show the theoretical designs of the Syn-CBS circuit with a weak M1-to-M2 link and strong M2-to-M1 link (A), or A strong M1-to-M2 link and weak M2-to-M1 link (B). The potential represents the stability of the steady states or the probabilities of the cells attracted to them.

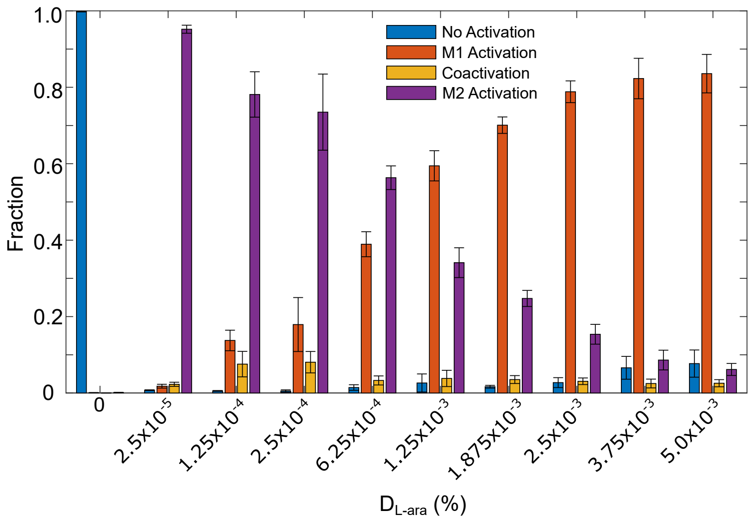


**Figure S2. The fraction of cells in different fates controlled by the one-strain Syn-CBS circuit with increasing the dose of L-ara ($D_{L\text{-}ara}$).** The fractions were estimated from flow cytometry data. Data displayed as mean ± s.d. (n = 4).

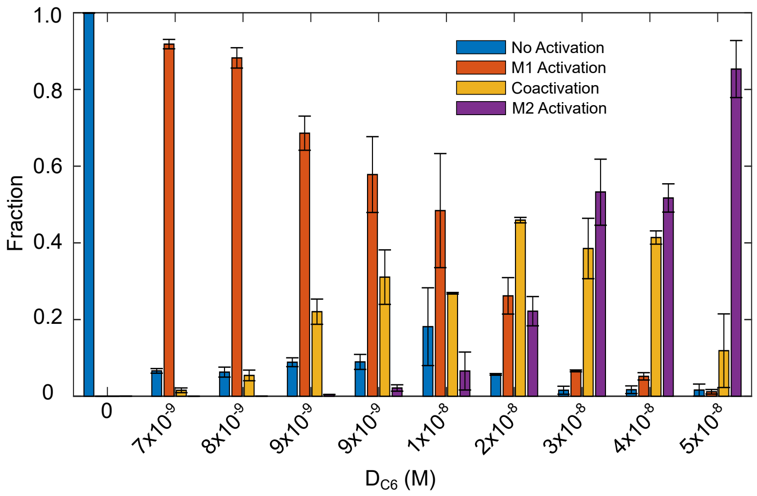


**Figure S3. The fraction of cells in different fates controlled by the two separated bistable switches system with increasing the dose of C6 ($D_{C6}$).** The fractions were estimated from flow cytometry data. Data displayed as mean ± s.d. (n = 3).

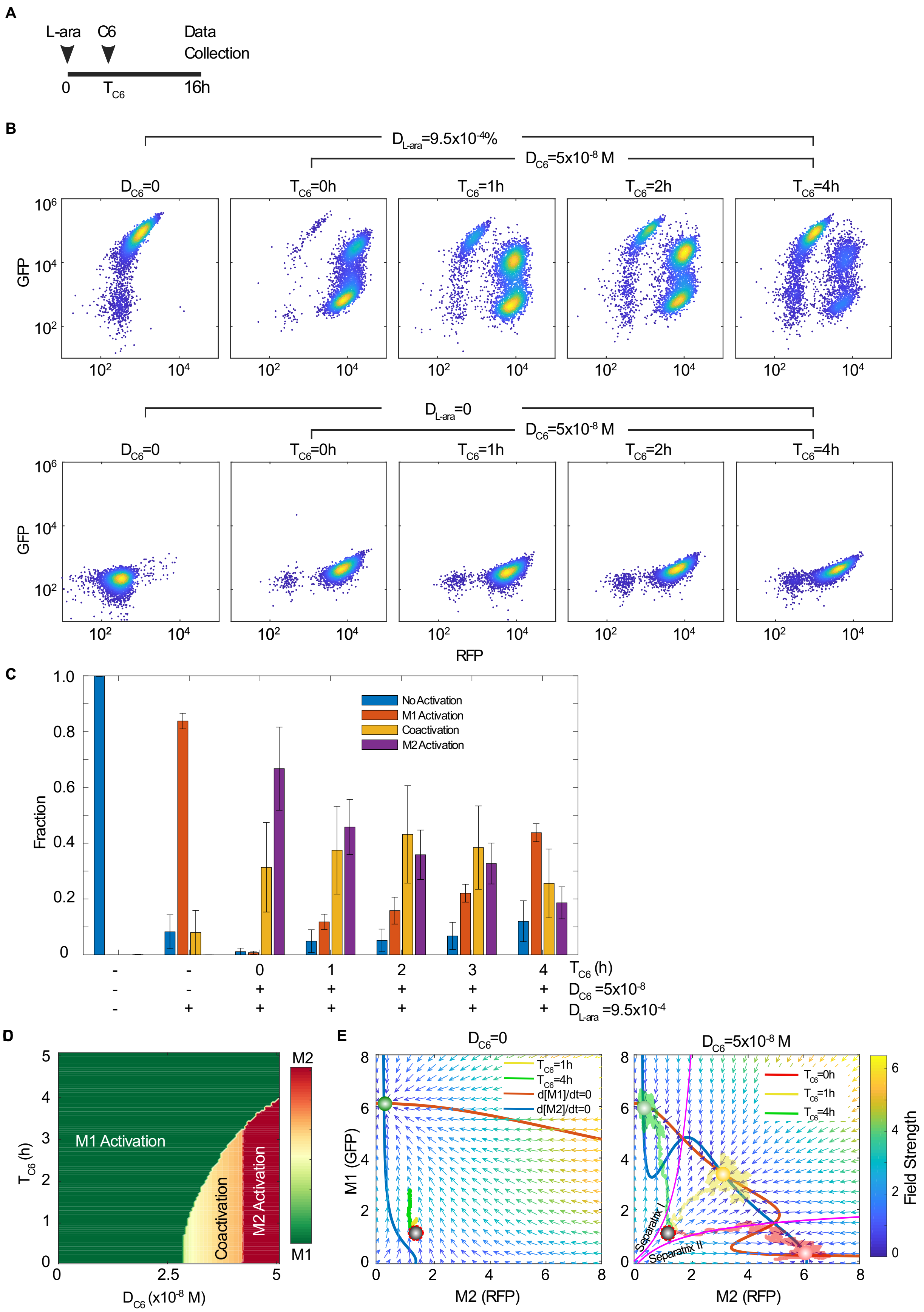
A
L-ara
C6
Data Collection
0
TC6
16h
B
DL-ara=9.5x10-4%
DC6=5x10-8 M
DC6=0
TC6=0h
TC6=1h
TC6=2h
TC6=4h
GFP
DL-ara=0
RFP
C
Fraction
No Activation
M1 Activation
Coactivation
M2 Activation
TC6 (h)
DC6 =5x10-8
DL-ara =9.5x10-4
D
M1 Activation
Coactivation
M2 Activation
TC6 (h)
DC6 (x10-8 M)
M2
M1
E
DC6=0
DC6=5x10-8 M
TC6=0h
TC6=1h
TC6=4h
d[M1]/dt=0
d[M2]/dt=0
M1 (GFP)
M2 (RFP)
Separatrix I
Separatrix II
Field Strength

**Figure S4. Resource competition between two separate bistable switches with sequential addition of the two inducers.** (A). Diagram of the experimental design. The doses of L-ara and C6 were fixed. L-ara was added at the time 0, and C6 was added at various time points. (B). Flow cytometry data shows cell state transitions with various dosing times of inducer C6 ($T_{C6}$) and a fixed dose of L-ara ($D_{L\text{-}ara}$) and C6 ($D_{C6}$). 10,000 events were recorded for each sample by flow cytometry. (C). The fraction of cells in different fates controlled by the two separated bistable switches system with various $T_{C6}$. The fractions were estimated from flow cytometry data. Data displayed as mean ± s.d. (n = 4). (D) Simulated cell fates in the space of the dose and timing of inducer C6. L-ara dose was fixed as $D_{L\text{-}ara}$=9.5x10$^{-4}$ %. (E). Simulated stochastic trajectories in the phase plane diagram. The initial state of the cells is set to the steady-state without any inducer (black circle). The nullclines of M1 and M2 are shown in red and blue, respectively, while separatrices are shown in pink. The three cell fates (red, green, and yellow circles) are found at the intersections of the two nullclines. The vector field of the system is represented by small arrows, where the color is proportional to the field strength. Three representative single-cell stochastic trajectories (green, yellow, and red trails) show the evolution of the system from the same initial condition ($D_{L\text{-}ara}$=0% and $D_{C6}$=0 M) to three different states with various $T_{C6}$. The dose of C6 was fixed as $D_{C6}$=0 M in the left panel and $D_{C6}$=5x10$^{-8}$ M in the right panel. L-ara was fixed as $D_{Lara}$=9.5x10$^{-4}$ %.

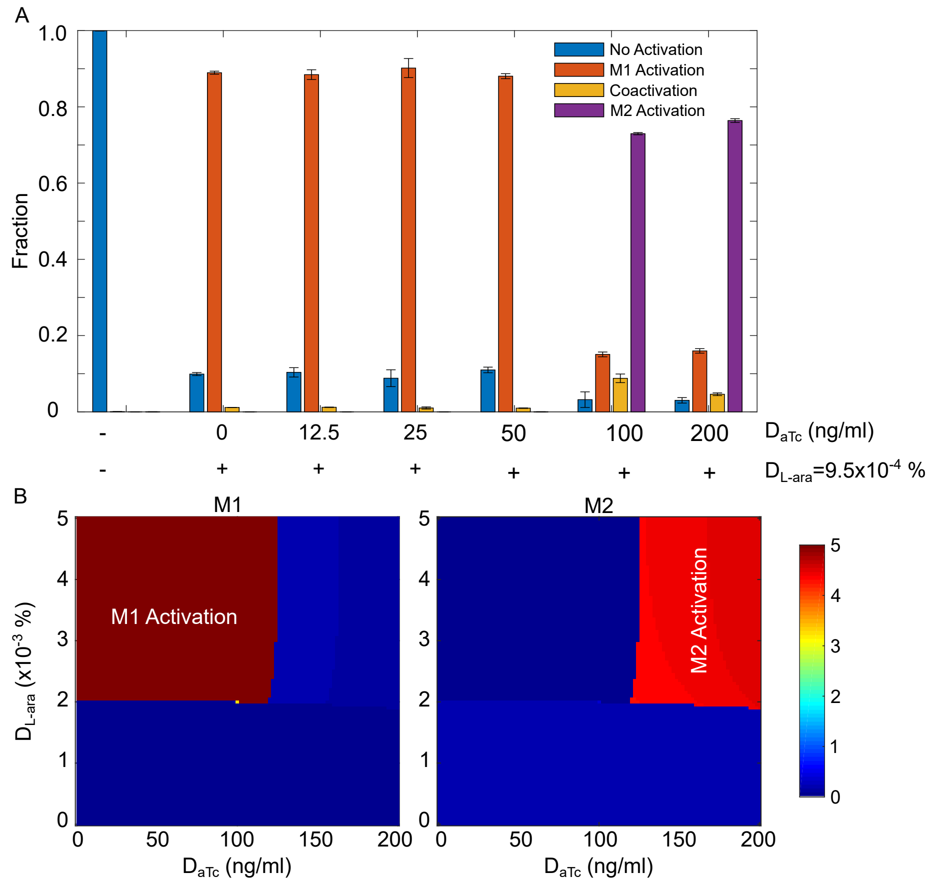


**Figure S5. Tune the outcomes of the resource competition by controlling relative strength of module connections.** (A) The fraction of cells in different fates controlled by the Syn-CBS circuit by increasing the dose of aTc ($D_{aTc}$) in the tetR module. The fractions were estimated from flow cytometry data. Data displayed as mean ± s.d. (n = 3). (B) Simulated cell fates in the doses space of two inducers L-ara and aTc.

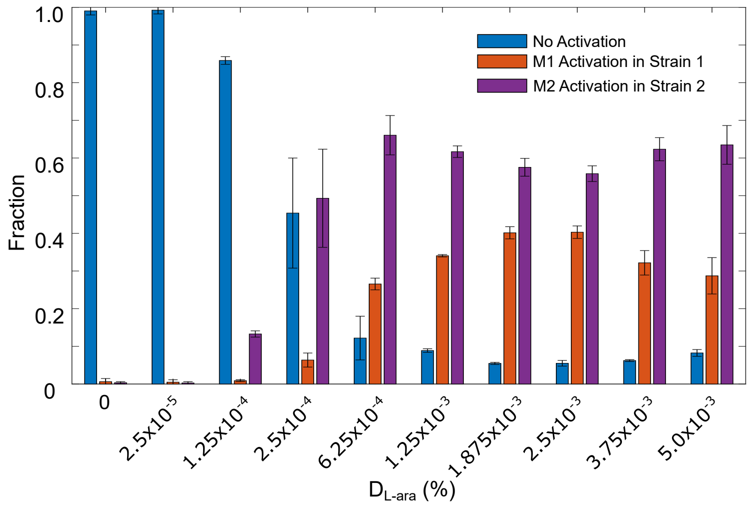


**Figure S6. The fraction of cells in different fates as a function of inducer L-ara controlled by the two-strain Syn-CSS circuits without the tetR module.** The fractions were estimated from flow cytometry data. Data displayed as mean ± s.d. (n = 3).

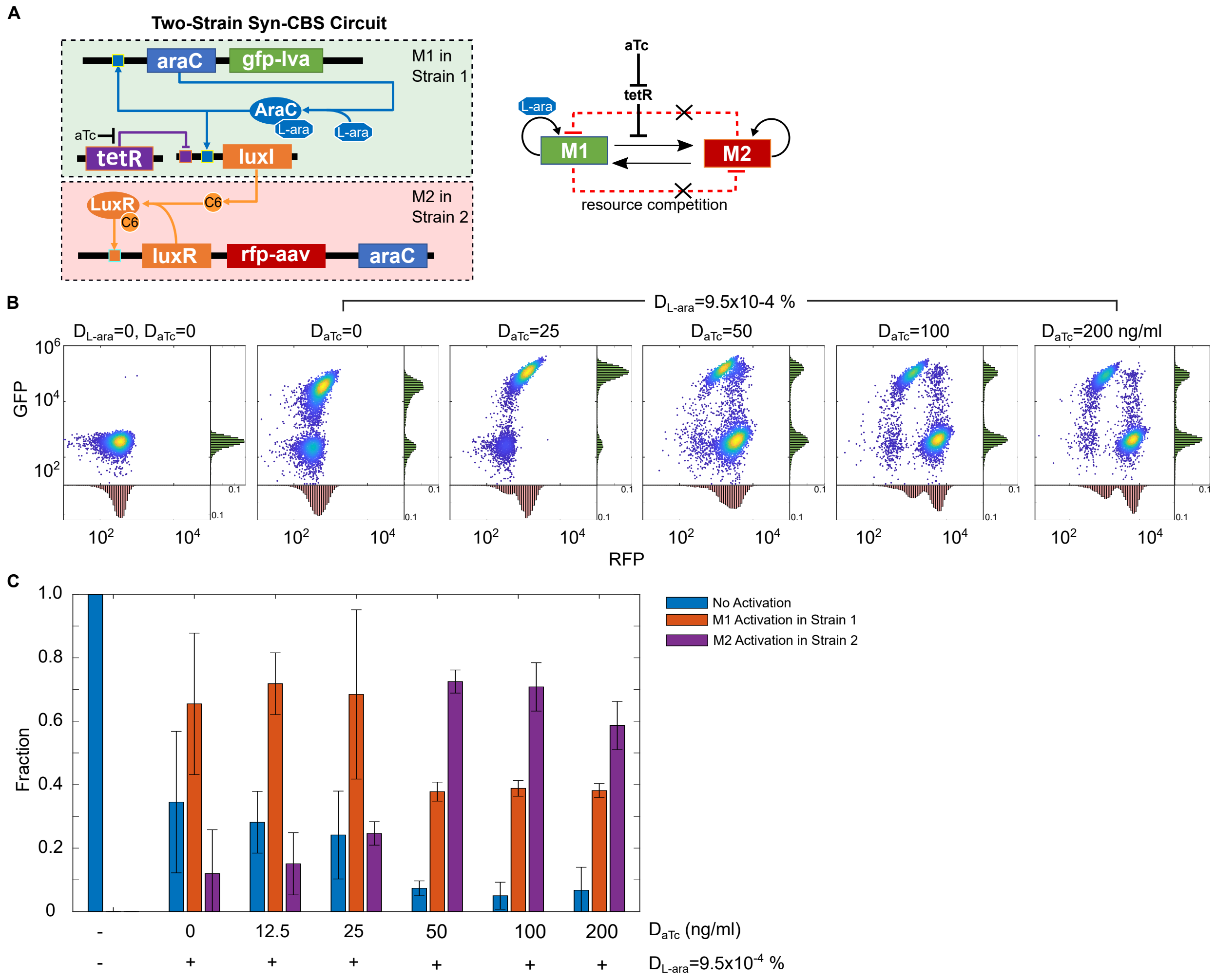


**Figure S7. Migration of resource competition with microbial consortia.** (A) Diagram of two-strain Syn-CBS circuits with a tetR module. (B) Flow cytometry data shows the expected stepwise cell state transitions by increasing the dose of aTc ($D_{aTc}$, ng/mL) in the two-strains Syn-CBS circuit with the tetR module. 10,000 events were recorded for each sample by flow cytometry. (C) The fraction of cells in different fates as functions of inducer aTc. The fractions were estimated from flow cytometry data. Data displayed as mean ± s.d. (n = 3).

**Table S1. BioBrick parts used in this paper.**

| BioBrick number | Abbreviation in the paper | Description |
|---|---|---|
| K206000 | Pbad | Inducible promoter activated by AraC and L-arabinose |
| C0061 | LuxI | 3oxo-C6-HSL producing enzyme |
| C0040 | TetR | Tetracycline repressor from transposon Tn10 |
| J23116 | Pcon | Constitutive promoter |
| J04031 | GFP | GFP generator |
| B0034 | RBS | Ribosome binding site |
| B0015 | Terminator | Transcriptional terminator (double direction) |
| PSB1C3 | PSB1C3 | High copy BioBrick backbone with chloramphenicol resistance |
| pSB3K3 | pSB3K3 | Medium copy BioBrick assembly backbone with kanamycin resistance |

**Table S2. List of monocistronical operons**

| ID | Description (promoter-gene) | sub-parts | Backbone |
|---|---|---|---|
| op9 | Pbad-araC | K206000+B0034+araC+B0010+B0012 | PSB1C3 |
| K750000 | Pbad-gfpLVA | K206000+B0034+K145915+B0010+B0012 | PSB1C3 |
| op12 | Pbad-luxRG2C | K206000+B0034+luxRG2C+B0010+B0012 | PSB1C3 |
| op97 | Plux-rfpAAV | Plux+B0034+rfpAAV+B0010+B0012 | PSB1C3 |
| op105 | Plux-luxRG2C | Plux+B0034+luxRG2C+B0010+B0012 | PSB1C3 |
| op101 | Plux-araC | Plux+B0034+araC+B0010+B0012 | PSB1C3 |
| op127 | J23116-tetR | J23116+B0034+P0440+B0010+B0012 | PSB1C3 |
| op111 | Pbad/tet-luxI | Pbad/tet+B0034+C0061+B0010+B0012 | PSB1C3 |
| op54 | Pbad-luxI | K206000+B0034+C0061+B0010+B0012 | PSB1C3 |

**Table S3. List of gene circuits**

| ID | Assembly from operons | Description | Backbone |
|---|---|---|---|
| CT61 | K750000+op9+op105+op97+op101+op54 | Pbad-GFPlva+Pbad-araC+Plux9-luxRG2C+Plux9-RFPaav+Plux9-araC+Pbad-luxI | **PSB3K3** |
| CT81 | op127+op111+K750000+op9+op105+op97+op101 | J23116-tetR+PBad/tet-luxI+Pbad-GFPlva+Pbad-araC+Plux9-luxRG2C+Plux9-RFPaav+Plux9-araC | **PSB3K3** |
| CT66 | op105+op97+op101 | Plux9-luxRG2C+Plux9-RFPaav+Plux9-araC | **PSB3K3** |
| CT67 | K750000+op9+op54 | PBADs-GFPlva+PBADs-araC+PBADs-LuxI | **PSB3K3** |
| CT82 | op127+K750000+op9+op111 | J23116-tetR+Pbad-GFPlva+Pbad-araC+Pbad/tet-luxI | **PSB3K3** |

**Mathematical modeling**

**Model for the Syn-CSB circuit without considering resource competition**

The Syn-CBS circuits are composed of two modules. In each module, there is one activator, which promotes the its own production, thus forming a self-activation motif. Specifically, in module 1 (M1), the AraC-L-ara dimer binds to promoter $P_{bad}$ to promote the production of itself, reporter GFP, and the signal C6 for module 2. In module 2 (M2), the LuxR-C6 dimer binds to the promoter $P_{lux}$ to induce the production of itself, reporter RFP, and another copy of araC. Thus, the two modules promote each other. The construction of the model for the AraC self-activation module is based on our previous works[1]. The LuxR self-activation module is similar to the AraC self-activation module and follows a similar equation. The connections of the two modules are mediated by the AraC-mediated production of C6 and the LuxR-mediated production of AraC. Here, for simplicity, we model the genes under the same promoter as one variable instead of modeling all the genes as separate variables. This simplification is reasonable given that the production rates for the genes under the same promoter should be similar as each operon constituting the circuits was constructed monocistronically. In this way, we can build a two-dimensional ordinary differential equations (ODEs) model with two variables, $M_1$ for the genes in modules 1 and $M_2$ for the gene in module 2. The level of C6 is based on M1 with one coefficient $\lambda_1$, and the total level of AraC includes the part in M1 and the part mediated by LuxR that is based on M2 with one coefficient $\lambda_2$. This two-dimensional ODE model allows us to do nullcline and direction field analysis directly. The mathematical model for the Syn-CBS circuit can be simplified by the following two equations:

$$\frac{dM_1}{dt} = \mathrm{f_1(M_1, M_2) - d_1 \cdot M_1}$$

$$\frac{dM_2}{dt} = \mathrm{f_2(M_1, M_2) - d_2 \cdot M_2}$$

Where $\mathrm{f_1} = \left(\mathrm{k_{01} + k_1 \cdot \frac{Sa \cdot AraC^2}{Sa \cdot AraC^2 + 1}}\right)$, $\mathrm{Sa = C_{min1} + (C_{max1} - C_{min1}) \cdot \frac{Lara^n}{Lara^n + J_1^n}}$ , $\mathrm{AraC = M_1 + M_2 \cdot \lambda_2}$,

$\mathrm{f_2} = \left(\mathrm{k_{02} + k_2 \cdot \frac{Su \cdot LuxR^2}{Su \cdot LuxR^2 + 1}}\right)$, $\mathrm{Su = C_{min2} + (C_{max2} - C_{min2}) \cdot \frac{C6^m}{C6^m + J_2^m}}$, , $\mathrm{C6 = M_1 \cdot \lambda_1}$, and $\mathrm{LuxR = M_2}$.

Here, $k_{01}$ and $k_{02}$ are the basic production rates of M1 and M2, while $k_1$ and $k_2$ are the maximum production rates of M1 and M2, respectively. $S_a$ describes how the production rate is regulated by inducer L-ara. $C_{max1}$ and $C_{min1}$ are the maximum and minimum affinities of the AraC dimers to the binding sites on the promoter $P_{bad}$. $S_u$ describes how the production rate is regulated by the LuxR. $C_{max2}$ and $C_{min2}$ are the maximum and minimum affinities of the LuxR dimers to the binding sites on the promoter $P_{lux}$. As well, $n$ represents the nonlinearity of the promoter activation by L-ara, and $d_1$ and $d_2$ are the degradation rates. The input of the system is the concentration of L-ara. The two reporters are $\mathrm{GFP} = \mathrm{M_1}$ and $\mathrm{RFP} = \mathrm{M_2}$. The model is suited to analyze the steady-state behavior of the system under conditions without resource competition. The theoretical analysis of the Syn-CBS circuit in Fig. 1 is based on this model. The fitted parameters are, unless otherwise mentioned: $\mathrm{C_{min1}} = 0.25$ , $\mathrm{C_{max1}} = 2$, $\mathrm{J_1} = 6 * 10^{-3}$, $n = 3$, $\mathrm{k_{01}} = 0.1$, $\mathrm{k_1} = 4$, $\mathrm{d_1} = 1$, $\mathrm{C_{min2}} = 0.2$, $\mathrm{C_{max2}} = 2$, $\mathrm{J_2} = 1.5$, $m = 3$, $\mathrm{k_{02}} = 0.1$, $\mathrm{k_2} = 5$, $\mathrm{d_2} = 1$, and $\mathrm{Lara} = 1.2 * 10^{-3}$. The strength of the connections between the two modules are set as $\lambda_1 = 0.25$ and $\lambda_2 = 0.2$ in Fig.1BC and $\lambda_1 = 0.5$ and $\lambda_2 = 0.03$ in Fig.1DE to demonstrate two possible theoretical designs of the synthetic cascading bistable switches.

**Model for the Syn-CSB circuit when considering resource competition**

The expectations from the mathematical model of the Syn-CSB circuit without resource competition are not consistent with the experimental data, thus we developed the general mathematical model for a synthetic gene circuit by considering the resources (RNA polymerase and ribosome) in the host cell (see the following section below) and applying it to the Syn-CBS circuit. The mathematical model for the Syn-CSB circuit is thus revised as follows:

$$\frac{dM_1}{dt} = (\mathrm{v_{01}} \cdot \mathrm{R_{01}} + \mathrm{v_1} \cdot \mathrm{R_1})/\mathrm{PF_Q} - \mathrm{d_1} \cdot \mathrm{M_1}$$

$$\frac{dM_2}{dt} = (\mathrm{v_{02}} \cdot \mathrm{R_{02}} + \mathrm{v_2} \cdot \mathrm{R_2})/\mathrm{PF_Q} - \mathrm{d_2} \cdot \mathrm{M_2}$$

where $\mathrm{R_1} = \frac{\mathrm{Sa \cdot AraC^2}}{\mathrm{Sa \cdot AraC^2 + 1}} \cdot N_{cp}$, $\mathrm{R_2} = \frac{\mathrm{Su \cdot LuxR^2}}{\mathrm{Su \cdot LuxR^2 + 1}} \cdot N_{cp}$, $\mathrm{Sa} = \mathrm{C_{min1}} + (\mathrm{C_{max1}} - \mathrm{C_{min1}}) \cdot \frac{\mathrm{Lara^n}}{\mathrm{Lara^n + J_1^n}}$, $\mathrm{Su} = \mathrm{C_{min2}} + (\mathrm{C_{max2}} - \mathrm{C_{min2}}) \cdot \frac{\mathrm{C6^m}}{\mathrm{C6^m + J_2^m}}$, $R_{01} = N_{cp}$, $R_{02} = N_{cp}$, $\mathrm{C6} = \mathrm{M_1} \cdot \lambda_1$, $\mathrm{AraC} = \mathrm{M_1} + \mathrm{M_2} \cdot \lambda_2$, $\mathrm{LuxR} = \mathrm{M_2}$, and $\mathrm{PF_Q} = (\frac{1}{\mathrm{Q_{01}}} + \frac{1}{\mathrm{Q_{02}}} + \frac{\mathrm{R_1}}{\mathrm{Q_1}} + \frac{\mathrm{R_2}}{\mathrm{Q_2}}) + 1$. Here, we now consider $N_{cp}$ as the copy number of the plasmid. The theoretical analysis of the Syn-CBS circuit in Fig. 2 and Fig. 4 is based on this model. The copy number of the plasmid is in a range of 20-30 in our system. Thus, we used $N_{cp} = 24$ in the mathematical model. The fitted parameters are, unless otherwise mentioned, $\mathrm{C_{min1}} = 0.003$, $\mathrm{C_{max1}} = 0.1275$, $\mathrm{J_1} = 0.75 * 10^{-3}$, $n = 3$, $\mathrm{v_{01}} = 0.0005$, $\mathrm{v_1} = 0.5$, $\mathrm{d_1} = 0.25$, $\mathrm{C_{min2}} = 0.005$, $\mathrm{C_{max2}} = 0.175$, $\mathrm{J_2} = 0.5$, $m = 3$, $\mathrm{v_{02}} = 0.0025$, $\mathrm{v_2} = 0.5$, $\mathrm{d_2} = 0.25$, $\lambda_1 = 1.25$, $\lambda_2 = 0.044$, $\mathrm{Q_{01}} = 300$, $\mathrm{Q_1} = 300$, $\mathrm{Q_{02}} = 3$, $\mathrm{Q_2} = 3$, and $\mathrm{Lara} = 0 \sim 5 * 10^{-3}$. L-ara is set to $1.25 * 10^{-3}$ in Fig. 2EF. The inducer aTc in the Syn-CBS circuit with the tetR module is set by the level of $\lambda_1$. aTc range is set to 0~200 by linearly scale $\lambda_1 = 0 \sim 1.25$ in Fig. S3.

**Mathematical model for the two separate switches system**

The two separate switches system was used to verify the resource competition between the two modules within the Syn-CBS circuit and the WTA behavior. Most parts of the system are the same as the Syn-CBS circuit except for the two links that connect the modules, including AarC-mediated production of C6 and LuxR-mediated production of AraC, were removed in two separate switches system. Thus, the mathematical model for the two separate switches system with resource competition is as follows:

$$\frac{dM_1}{dt} = (\mathrm{v}_{01} \cdot \mathrm{R}_{01} + \mathrm{v}_1 \cdot \mathrm{R}_1)/\mathrm{PF_Q} - \mathrm{d}_1 \cdot \mathrm{M}_1$$

$$\frac{dM_2}{dt} = (\mathrm{v}_{02} \cdot \mathrm{R}_{02} + \mathrm{v}_2 \cdot \mathrm{R}_2)/\mathrm{PF_Q} - \mathrm{d}_2 \cdot \mathrm{M}_2$$

where$\mathrm{R}_1 = \frac{\mathrm{Sa \cdot AraC^2}}{\mathrm{Sa \cdot AraC^2+1}} \cdot N_{cp}$ , $\mathrm{R}_2 = \frac{\mathrm{Su \cdot LuxR^2}}{\mathrm{Su \cdot LuxR^2+1}} \cdot N_{cp}$, $\mathrm{Sa} = \mathrm{C_{min1}} + (\mathrm{C_{max1}} - \mathrm{C_{min1}}) \cdot \frac{\mathrm{L0^n}}{\mathrm{L0^n+J1^n}}$, $\mathrm{Su} = \mathrm{C_{min2}} + (\mathrm{C_{max2}} - \mathrm{C_{min2}}) \cdot \frac{\mathrm{C6^m}}{\mathrm{C6^m+J2^m}}$, $R_{01} = N_{cp}$, $R_{02} = N_{cp}$, $\mathrm{AraC} = \mathrm{M}_1$, $\mathrm{LuxR} = \mathrm{M}_2$, and $\mathrm{PF_Q} = (\frac{1}{\mathrm{Q}_{01}} + \frac{1}{\mathrm{Q}_{02}} + \frac{\mathrm{R}_1}{\mathrm{Q}_1} + \frac{\mathrm{R}_2}{\mathrm{Q}_2}) + 1$. The inputs to this system are L-ara and C6, which control the M1 switch and M2 switch separately. The theoretical analysis of the Syn-CBS circuit in Fig. 3 and Fig. S3 is based on this model. The parameters are, unless otherwise mentioned, $\mathrm{C_{min1}} = 0.003$, $\mathrm{C_{max1}} = 0.1275$, $\mathrm{J}_1 = 4.38 * 10^{-4}$, $n = 3$, $\mathrm{v}_{01} = 0.0125$, $\mathrm{v}_1 = 0.5$,$\mathrm{d}_1 = 0.25$, $\mathrm{C_{min2}} = 0.005$, $\mathrm{C_{max2}} = 0.175$, $\mathrm{J}_2 = 2.5 * 10^{-8}$, $m = 3$, $\mathrm{v}_{02} = 0.0125$, $\mathrm{v}_2 = 0.5$, $\mathrm{d}_2 = 0.25$, $\mathrm{Q}_{01} = 300$, $\mathrm{Q}_1 = 300$, $\mathrm{Q}_{02} = 3.6$, $\mathrm{Q}_2 = 3.6$, $\mathrm{Lara} = 0 \sim 10 * 10^{-4}$, and $C6 = 0 \sim 5 * 10^{-8}$. L-ara is set to $9.5 * 10^{-4}$ and C6 is set to $3 * 10^{-8}$ in Fig. 3D.

**General model for the synthetic circuit with resource competition**

For a synthetic circuit with multiple genes, we first considered the general model without resource competition. The ordinary differential equations of mRNA and protein products for each gene follow:

$$\frac{dmRNA_i}{dt} = k_{mi} \cdot R_i - d_{m_i} \cdot mRNA_i$$

$$\frac{dP_i}{dt} = k_{pi} \cdot mRNA_i - d_{Pi} \cdot P_i$$

where $R_i$ is the number of active promoters for each gene that is bound by transcription factors ($DNA_i$:$TF$). For this model, the resources such as RNAP and ribosome are not considered yet.

We then considered the transcription resource RNAP in the model. To do so, we needed to consider the binding/unbinding of the RNAP to the active promoter $DNA{:}TF$ in order to start transcription,

$$DNA_i{:}TF + RNAP \xleftrightarrow{k_f^{RNAP},k_r^{RNAP}} DNA_i{:}TF{:}RNAP \xrightarrow{k'_m} mRNA_i + RNAP{:}TF$$

Here, we assume that the concentrations of the RNA polymerases are constant [2]. It is noted that all the promoters in the synthetic gene circuit compete for the available RNAP within the host cell. Thus, the transcription rate for each gene follows the Michaelis–Menten kinetics with competitive inhibition by all other genes:

$$k'_m \cdot RNAP_t \cdot \frac{R_i/J_{m_i}}{\sum R_i/J_{m_i} \quad +1} = \frac{k'_{mi} \cdot RNAP_t}{J_{mi}} \cdot \frac{R_i}{\sum R_i/J_{m_i} \quad +1} = k_{mi} \cdot \frac{R_i}{\sum R_i/J_{m_i} \quad +1}$$

where $k_{mi} = \frac{k'_{mi} \cdot RNAP_t}{J_{m_i}}$, $J_m = \frac{k'_m + k_r^{RNAP}}{k_f^{RNAP}}$, $RNAP_t$ is the total available RNAP in the host that can be used for the synthetic gene circuits, and $J_{m_i}$ is the Michaelis constant. At $DNA{:}TF_n = J_{m_i}$, the transcription rate is at half-maximum transcription rate when RNAP is saturated.

Thus, the ODE of mRNA is revised as:

$$\frac{dmRNA_i}{dt} = k_{mi} \cdot \frac{R_i}{\sum R_i/J_{m_i} \quad +1} - d_{mi} \cdot mRNA_i$$

which can be further simplified to

$$\frac{dmRNA_i}{dt} = \frac{k_{mi} \cdot R_i}{PF_m} - d_{mi} \cdot mRNA_i \, .$$

Here, $PF_m = \sum R_i/J_{m_i} + 1$. Under the condition $R/J_m \ll 1$ (i.e., $RNAP$ is far from saturated), $PF_m = \sum R_i/J_{m_i} + 1$. The ODE of mRNA is the same as the one without RNAP competition (Eq. 1).

We further consider the competition of translation resources such as ribosome. To do so, we needed to consider the binding/unbinding of the ribosome to each mRNA in order to start translation,

$$mRNA_i + Ribosome \xleftrightarrow{k_f^{Ribosome},k_r^{Ribosome}} mRNA_i{:}Ribosome \xrightarrow{k'_p} P_i + mRNA_i$$

Here the total concentrations of ribosomes is considered to be constant [2]. All the mRNAs compete for the available ribosome. Thus, the translation rate for each mRNA also follows the Michaelis–Menten kinetics with competitive inhibition by all other mRNAs,

The translation rate of $mRNA_i$ is

$$k'_P Ribosome_t \frac{k_{pi} \cdot mRNA_i}{\sum mRNA_i/J_{p_i} + 1} = k'_P \cdot Ribosome_t \cdot \frac{mRNA/J_p}{mRNA/J_p + 1} = \frac{k_{p_i} \cdot mRNA_i}{\sum mRNA_i/J_{p_i} + 1}$$

where $k_{p_i} = \frac{k'_P \cdot Ribosome_t}{J_{p_i}}$, $J_{p_i} = \frac{k'_P + k_r^{Ribosome}}{k_f^{Ribosome}}$, $Ribosome_t$ is the total available ribosome in the host which can be used for the synthetic gene circuits, and $J_{p_i}$ is the Michaelis constant. At $mRNA_i = J_{p_i}$, the transcription rate is at half-maximum transcription rate when the ribosome is saturated.

Thus, the ODE of each protein product is:

$$\frac{dP_i}{dt} = k_{pi} \frac{mRNA_i}{\sum mRNA_i/J_{p_i} + 1} - d_{Pi} \cdot P_i$$

which can be further simplified to

$$\frac{dP_i}{dt} = \frac{k_{pi} \cdot mRNA_i}{\mathrm{PF_p}} - d_{Pi} \cdot P_i. \quad (8)$$

Here, $\mathrm{PF_p} = \sum mRNA_i/J_{p_i} + 1$. Under the condition $mRNA/J_p \ll 1$ (i.e., ribosome is far from saturated), $\mathrm{PF_p} = 1$, and the ODE of mRNA is the same as the one without RNAP competition (Eq. 2).

We further simplified equations by elevating the equations of miRNAs

$$\frac{dP}{dt} = \frac{k_{pi} \cdot k_{mi}/d_{mi} \cdot R_i}{PF_m * \mathrm{PF_p}} - d_{Pi} * P_i = \frac{v_{pi} \cdot R_i}{PF_m \cdot \mathrm{PF_p}} - d_{Pi} \cdot P_i \quad (10)$$

Where $v_{pi}(= k_{pi} \cdot \frac{k_{mi}}{d_{m_i}})$ is a lumped parameter that represents the overall gene expression rate. $\mathrm{PF_p} = \sum \frac{k_{mi}/d_{mi} \cdot R_i}{PF_m} \frac{1}{J_{p_i}} + 1 = \sum \frac{R_i/L_i}{PF_m} + 1$, $L_i = \frac{J_{p_i} d_{mi}}{k_{mi}} = \frac{d_{mi}\left(k_{r_i}^{ribosome} + k_{p_i}\right)}{k_{mi} k_{f_i}^{ribosome}}$, and $PF_m = \sum R_i/J_{m_i} + 1$. Thus, $PF_m \cdot \mathrm{PF_p} = \sum \frac{R_i}{J_{m_i}} + \sum \frac{R_i}{L_i} + 1 = \sum R_i/Q_i + 1$.

Thus, the final simplified general model for the synesthetic gene circuit with resource competition is

$$\frac{dP_i}{dt} = \frac{v_{pi} \cdot R_i}{\mathrm{PF_Q}} - d_{Pi} \cdot P_i \quad (11)$$

where $\mathrm{PF_Q} = \sum R_i/Q_i + 1$, and the new lumped parameter $Q_i = \frac{1}{\frac{1}{J_{mi}}+\frac{1}{L_i}}$ indicates the overall capacity of limited resources in the host cell for synthetic gene circuits. While the ribosome is the main limited resource for synthetic gene circuits [2-5], the contribution of the translational capacity to the lumped parameter Q is more significant than the transcriptional capacity.

**Stochastic Models**

We also developed stochastic models for all of the synthetic circuits with or without resource competition, which generally can be described as birth-and-death stochastic processes that governed the production and degradation rates in the ODE models. A system size factor Ω is introduced to convert the concentration of each variable X (i.e., x = [x]·Ω). The stochastic transition processes and the corresponding propensity function for all the models are described in Table S4. Gillespie algorithm was used for the stochastic simulation.

**Potential landscape computation**

For a general two-dimensional system described with the following ordinary differential equations

$$\frac{d[X]}{dt} = f_1([X],[Y]) - g_1([X],[Y]),$$
$$\frac{d[Y]}{dt} = f_2([X],[Y]) - g_2([X],[Y]),$$

where $[X],[Y]$ are the concentration of the two variables, and both $f_i([X],[Y])$ and $g_i([X],[Y])$ represent the production and degradation rates for each variable, respectively.

The corresponding Chemical Master equation (CME)[6] is:

$$\frac{d\,P(X,Y,t)}{dt} = f_1(X-1,Y)P(X-1,Y) + g_1(X+1,Y)P(X+1,Y) + f_2(X,Y-1)P(X,Y-1) + g_2(X,Y+1)P(X,Y+1) - \big(f_1(X,Y) + g_1(X,Y) + f_2(X,Y) + g_2(X,Y)\big)P(X,Y),$$

where $X,Y$ are the number of molecules, and $P(X,Y,t)$ represents the probability of the system in state $(X,Y)$ at time $t$. The steady-state distribution $P_{ss}$ can be obtained by solving the following equation:

$$0 = f_1(X-1,Y)P_{ss}(X-1,Y) + g_1(X+1,Y)P_{ss}(X+1,Y) + f_2(X,Y-1)P_{ss}(X,Y-1) + g_2(X,Y+1)P_{ss}(X,Y+1) - \big(f_1(X,Y) + g_1(X,Y) + f_2(X,Y) + g_2(X,Y)\big)P_{ss}(X,Y)$$

To numerically solve for the $P_{ss}$, we rewrote the above equation in matrix form:

$$A \cdot P_{ss} = 0.$$

where A is the transition rate matrix from state $(X+i, Y+j)$ to state $(X, Y)$, defined as

$$A\,(X+i, Y+j \to X, Y) = \begin{cases} -\big(f_1(X,Y) + g_1(X,Y) + f_2(X,Y) + g_2(X,Y)\big) & (i=0, j=0) \\ f_1(X-1,Y) & (i=-1, j=0) \\ g_1(X+1,Y) & (i=\ \ 1, j=0) \\ f_2(X,Y-1) & (i=0, j=-1) \\ g_2(X,Y+1) & (i=0, j=\ \ 1) \\ 0 & otherwise \end{cases}$$

No-flux boundary conditions were used to conserve probability. By solving the above linear equation with the Gauss-Seidel method, we found the steady-state distribution $P_{ss}$ and estimated the potential landscape $U \approx -\ln(P_{ss})$ [7].

**Table S4. Stochastic models for the synthetic gene circuits.**

| Reaction | Description | Propensity function |
|---|---|---|
| Φ→M1 | Basal production rate of M1 | $(v_{01} \cdot R_{01})/PF_Q \cdot \Omega$ |
| Φ→M1 | Production rate of M1 | $(v_1 \cdot R_1)/PF_Q \cdot \Omega$ |
| M1→Φ | Degradation rate of M1 | $d_1 \cdot M1$ |
| Φ→M2 | Basal production rate of M2 | $(v_{02} \cdot R_{02})/PF_Q \cdot \Omega$ |
| Φ→M2 | Production rate of M2 | $(v_2 \cdot R_2)/PF_Q \cdot \Omega$ |
| M2 →Φ | Degradation rate of M2 | $d_2 \cdot M2$ |

For the Syn-CBS circuit with resource competition:

$$R_1 = \frac{Sa \cdot AraC^2}{Sa \cdot AraC^2 + \Omega^2} \cdot N_{cp}, \quad R_2 = \frac{Su \cdot LuxR^2}{Su \cdot LuxR^2 + \Omega^2} \cdot N_{cp}, \quad Sa = C_{min1} + (C_{max1} - C_{min1}) \cdot \frac{L0^n}{L0^n + J1^n},$$

$$Su = C_{min2} + (C_{max2} - C_{min2}) \cdot \frac{C6^m}{C6^m + J2^m}, \quad R_{01} = N_{cp}, \quad R_{02} = N_{cp}, \quad AraC = M_1 + M_2 \cdot \lambda_2,$$

$$C6 = M_1 \cdot \lambda_1, LuxR = M_2, PF_Q = (\frac{1}{Q_{01}} + \frac{1}{Q_{02}} + \frac{R_1}{Q_1} + \frac{R_2}{Q_2}) + 1.$$

For the two separate switches system with resource competition:

$$R_1 = \frac{Sa \cdot AraC^2}{Sa \cdot AraC^2 + \Omega^2} \cdot N_{cp}, \quad R_2 = \frac{Su \cdot LuxR^2}{Su \cdot LuxR^2 + \Omega^2} \cdot N_{cp}, \quad Sa = C_{min1} + (C_{max1} - C_{min1}) \cdot \frac{L0^n}{L0^n + J1^n},$$

$$Su = C_{min2} + (C_{max2} - C_{min2}) \cdot \frac{C6^m}{C6^m + J2^m}, \quad R_{01} = N_{cp}, \quad R_{02} = N_{cp}, \quad AraC = M_1,$$

$$LuxR = M_2, \quad PF_Q = \left(\frac{1}{Q_{01}} + \frac{1}{Q_{02}} + \frac{R_1}{Q_1} + \frac{R_2}{Q_2}\right) + 1.$$
